# Relative Mutant N501Y SARS-CoV-2 Spike Protein RBD Inhibition of Anti-Spike Protein IgG and ACE-2 Binding to Spike Protein Species

**DOI:** 10.1101/2021.04.26.441517

**Authors:** Melvin E. Klegerman, Jeffrey D. Cirillo, David D. McPherson

## Abstract

In the SARS-CoV-2 coronavirus pandemic of 2019 (COVID-19), it has become evident that the ACE-2 receptor-binding domain (RBD) of the viral spike protein (SP) is the target of neutralizing antibodies that comprise a critical element of protective immunity to the virus. The most definitive confirmation of this contention is that the two mRNA COVID-19 vaccines in general use, which elicit antibodies specific for the RBD, exhibit approximately 95% protective efficacy against COVID-19. A potential challenge to vaccine efficacy is the emergence of SARS-CoV-2 variants possessing multiple mutations affecting amino acid residues in the RBD. Of concern are variants that arose in the United Kingdom, Brazil and South Africa. One of the variants, designated B.1.351, has shown a higher transmissibility due to greater affinity for the ACE-2 receptor and decreased neutralization by convalescent plasma, therapeutic monoclonal antibodies, and post-vaccination plasma. Common to several of the variants is the N501Y mutation in the RBD, which may be responsible for at least part of the observed variant properties. To test this hypothesis, we measured the ability of the Y501 RBD to inhibit binding of the wild type RBD and full SP (S1 + S2) to the ACE-2 protein and a human monoclonal IgG antibody elicited to the wild type RBD, relative to the wild type RBD in two enzyme-linked immunosorbent assays (ELISAs). We found no significant difference in the IC_50_ of the two RBD species’ inhibition of ACE-2 binding, but unexpectedly found that the IC_50_ of the wild type RBD inhibition of antibody binding was nearly twice that of the Y501 RBD, reflecting a lower affinity. These results suggest that the individual N501Y mutation does not contribute to altered viral properties by itself, but may contribute to a collective conformational shift produced by multiple mutations.

## INTRODUCTION

The coronavirus disease outbreak of 2019 (COVID-19), caused by a newly emerging coronavirus, severe acute respiratory syndrome coronavirus 2 (SARS-CoV-2), has infected more than 110 million people worldwide, resulting in more than 2.4 million deaths (500,000 in the U.S.), little more than 1 year after initial cases appeared in Wuhan, China, in December 2019. It has now become apparent that neutralizing antibodies specific for the viral spike protein receptor-binding domain (RBD), which binds to the angiotensin converting enzyme-2 (ACE-2) receptor on host cells, comprise the critical element of protective immunity to the virus (2). This contention is supported by the finding that the RBD is immunodominant and the target of 90% of the neutralizing activity present in SARS-CoV-2 immune sera (3). The BNT162b2 mRNA and mRNA-1273 SARS-CoV-2 vaccines, both coding for the spike glycoprotein, developed by Pfizer-BioNTech and Moderna, respectively, elicit antibodies specific for the RBD and exhibit approximately 95% protective efficacy against SARS-CoV-2 infection (4-6).

Several mutant SARS-CoV-2 strains have raised concerns regarding increased transmissibility, lethality, and escape from extant immunity, relative to the predominant pandemic strain, D614G, itself a mutant that arose soon after viral spread from Wuhan, China. Most notable of these mutant strains are B.1.1.7, B.1.351 and P.1, which arose in the U.K., South Africa and Brazil, respectively, during late 2020 to early 2021(7-11). The South African variant is of concern, with 21 mutations, including 3 in the RBD, since it has been shown to exhibit reduced neutralization titers, not only against therapeutic monoclonal antibodies, but also COVID-19 convalescent and post-vaccination plasma samples (10-13). Recently, it has been confirmed that these reductions of neutralizing antibody function were sufficient to impair vaccine efficacy appreciably (14).

Common to all of the three major variant strains is the RBD mutation N501Y, which signifies a substitution of L-tyrosine for L-asparagine at residue 501 of the spike protein. It has been proposed that a stronger hydrogen bond formed between the mutant residue and the Tyr 41 of the human ACE-2 molecule is responsible for the greater transmissibility of the viral variants (15-18). A laboratory strain of SARS-CoV-2 bearing the Y501 residue, however, did not exhibit decreased neutralizing capacity relative to the N501 strain by 20 post-BNT162b2 vaccination sera (6). We have utilized RBD inhibition of an enzyme-linked immunosorbent assay (ELISA) developed to measure binding of human IgG antibodies to the SARS-CoV-2 spike protein (SP) to test the hypotheses that the N501Y mutation specifically exhibits stronger SP binding to the ACE-2 receptor protein and weaker binding to a monoclonal human antibody raised against the native SP RBD.

## METHODS

### ELISA for Human IgG Antibodies Specific for the SARS-CoV-2 Spike Protein

This is a direct ELISA in which the antigen is adsorbed directly onto microtiter wells, as we previously described for fibrinogen (19). Incubation volumes were 50 µl and all incubations except for the initial coating step were at 37° C. After the blocking step, all incubations were followed by three washes with 0.02 M phosphate-buffered saline, pH 7.4 with 0.05% Tween-20 (PBS-T). Test wells were coated with 5 µg recombinant spike protein (rSP; S1+S2; R&D Systems, Minneapolis, MN)/ml coating buffer (0.05 M sodium bicarbonate, pH 9.6) overnight at 4° C. Well contents were aspirated and all wells (including background wells for each sample and standard dilution) were blocked for 1 hour with conjugate buffer (1% bovine serum albumin in 0.05 M Tris, pH 8.0, with 0.02% sodium azide). Human anti-SP IgG standards (chimera, GenScript, Piscataway, NJ) or human ACE-2 Fc (chimera, R&D Systems) in serial dilutions of 500-31.3 ng/ml PBS-T were incubated (n = 6) for 2 hours. All wells were then incubated with 3,000-fold diluted goat anti-human IgG-alkaline phosphatase conjugate (Sigma-Aldrich, St. Louis, MO) for 1 hour. The assay was developed by adding substrate buffer (0.05 M glycine buffer, pH 10.5, with 1.5 mM magnesium chloride) to each well, followed by 4 mg paranitrophenylphosphate (PNPP)/ml substrate buffer (for a total volume of 100 µl/well) and incubating for 15 minutes. The reaction was terminated by adding 50 µl 1 M sodium hydroxide to each well. Plates were read at 405 nm wavelength with a BioTek ELx808 multiwell plate reader.

The OD of background wells was subtracted from test well ODs. The net OD of antibody standard wells was plotted vs. IgG antibody concentration, which obeys a hyperbolic relation. For determination of unknown sample antibody concentrations, the curve fit equation was solved for x. Binding affinities with correction for conjugate incubation perturbation were determined as previously described (19). A correction nomogram for this ELISA is reproduced in Figure 1A.

**Figure 1.**
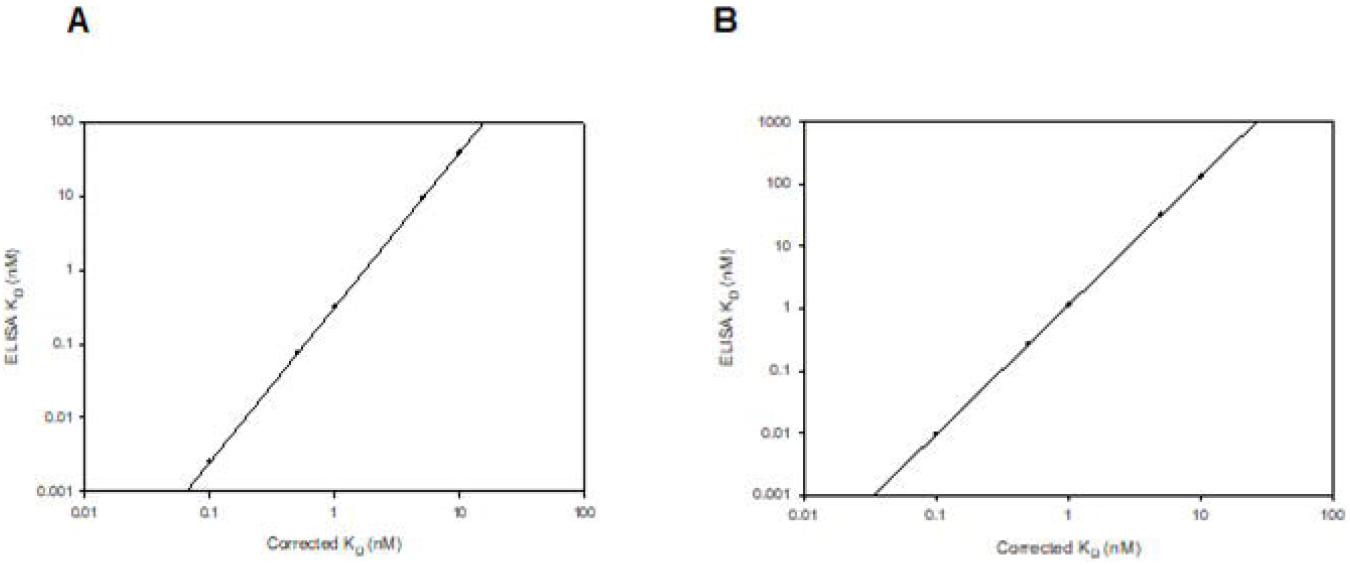
Nomograms for antibody binding affinity correction for equilibrium permutations following antibody-antigen incubation based on Underwood (1). Each point represents the calculated apparent K_D_ for a given “true” K_D_. A. rSP ELISA; log ELISA K_D_ = 2.098 log Corrected K_D_ - 0.502. B. RBD ELISA; log ELISA K_D_ = 2.075 log Corrected K_D_ + 0.060.

### ELISA for Human IgG Antibodies Specific for the SARS-CoV-2 Spike Protein Receptor-Binding Domain (RBD)

This is a sandwich ELISA in which a capture antibody, rabbit anti-mouse IgG, is adsorbed onto microtititer wells, followed by antigen capture. Test wells were coated with 2,000X diluted rabbit anti-mouse IgG (Sigma-Aldrich) in coating buffer overnight at 4° C. After blocking, 0.5 µg RBD-mFc (GenScript)/ml PBS-T was added to all wells and incubated for 2 hours. Human anti-SP IgG standards (in serial dilutions of 500-31.3 ng/ml PBS-T; n = 6) or human ACE-2 Fc (200-25 ng/ml; n = 3) were incubated for 1 hour. The rest of the ELISA protocol was the same as the rSP ELISA. The ODs of PBS-T only wells run in each assay were subtracted from test well ODs. Calculations of standards and sample antibody concentrations, as well as binding affinities, were performed as for the rSP ELISA. A correction nomogram for this ELISA is reproduced in Figure 1B.

### ELISA Inhibition by RBD Variants

Human anti-SP IgG or ACE-2 Fc was mixed with recombinant SARS-CoV-2 spike RBD His tag (R&D Systems) or recombinant SARS-CoV-2 spike RBD (N501Y)-His tag (Sino Biological, Wayne, PA), both comprising R319-F541 (MW 26 kDa), to produce serial dilutions of 0-500 ng/ml of RBD in the presence of 25 ng/ml of antibody standard for primary incubation following the blocking step of the rSP ELISA. Serial dilutions of 0-250 ng/ml of RBD in the presence of 250 ng/ml of ACE-2 Fc were utilized for secondary incubation of the RBD ELISA. IC_50_ values of inhibition curves were determined by exponential decay regression analysis using SigmaPlot 10 software, as (−0.693)/(-k), where k is the decay constant, expressed as molar concentration.

### Statistics

All data are expressed as mean ± standard deviation (SD). Significances, especially p < 0.05, were identified as a difference of calculated values. Differences between mean OD values of inhibition curves (p ≤ 0.05) were determined by normal variance analysis (t-test).

## RESULTS

Composite standard curves for the rSP and RBD ELISAs are shown in Figure 2. Antibody (Ab) standard binding affinity for the spike protein, with correction for post-antibody ELISA incubations (19), was found to be 1.37 ± 0.47 nM (K_D_)(SD, n = 21). Binding affinity of ACE-2 Fc for the spike protein was found to be 2.16 ± 0.67 nM (n =5), which is similar to published values (2). The corresponding values for the RBD ELISA were 0.55 ± 0.17 nM (n = 17) and 0.72 ± 0.21 nM (n = 3), respectively. Recoveries of IgG antibody standard from pooled normal plasma ranged from 94.5% to 132.5% (mean = 109.7%) from 6.25-100 ng Ab/ml.

**Figure 2.**
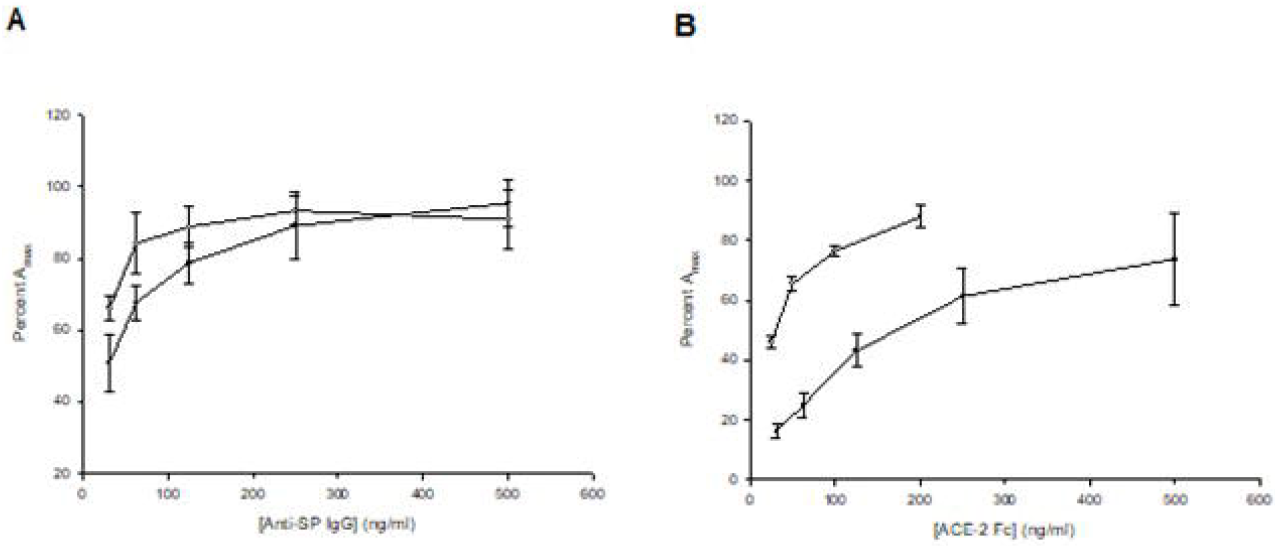
Composite dose-response curves for anti-spike protein lgG standards (A) and ACE-2 human Fc chimera protein (B) in the rSP (solid circles) and RBD (open circles) ELISAs. Points are the means of 6 determinations, except for the ACE-2 Fc RBD curve (n = 3); bars= SD.

Inhibition curves of ACE-2 Fc binding to the wild type RBD-mFc by the wild type RBD and the N501Y RBD were virtually identical (Fig. 3A). IC_50_ values were 2.32 nM and 2.52 nM, respectively. The former value was not different from rSP-ACE-2 Fc binding affinity determined by ELISA. Inhibition curves of the anti-SP IgG chimera standard binding to the wild type spike protein (S1 + S2) in the rSP ELISA by the wild type RBD and the N501Y RBD (Fig. 3B) were parallel, but different (all points, p < 0.05). The IC_50_ of the former was 20.8 nM, while that of the latter was 13.1 nM, indicating that the affinity of antibody binding to the N501Y RBD was actually nearly twice as great as to the wild type RBD.

**Figure 3.**
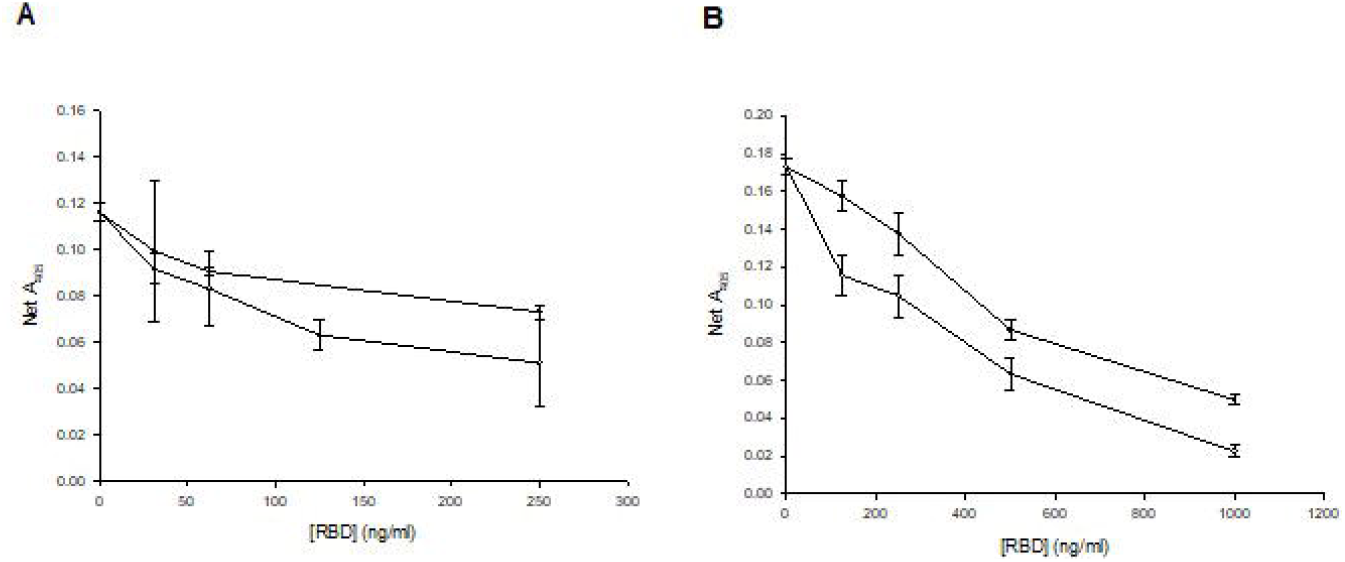
Inhibition curves of RBD species mixed with primary ligands in ELISAs. Filled circles: wild type RBD; open circles: N501Y RBD. A) ACE-2 Fc binding to RBD-mFc in RBD ELISA. B) Anti-SP lgG chimera binding to rSP in spike protein ELISA. Points are the mean of 3 determinations; bars = SD.

## DISCUSSION

These data demonstrate that a) the Y501 spike protein does not have greater affinity for the ACE-2 receptor protein than the N501 SP; b) the Y501 spike protein may have an even greater affinity for a human monoclonal antibody specific for the wild type SP than the N501 SP; and c) recombinant RBD inhibition of ligand binding to immobilized spike protein or RBD is a valid approach to assessing the relevant molecular impact of viral mutations.

Although there is ample evidence that the B.1.351 SARS-CoV-2 variant is more transmissible and less reactive with convalescent, post-vaccination, and therapeutic monoclonal neutralizing IgG antibodies than the predominant D614G pandemic strain, it is not at all clear what role the N501Y mutation plays in that shift. Most of the studies showing impaired antibody reactivity involved neutralization of whole virus infectivity *in vitro*. Neutralization of pseudoviruses engineered to express only the N501 or Y501 residues by 20 post-BNT162b2 vaccination sera was equivalent (6). Pseudoviruses engineered to express the K417N, E484K, N501Y and D614G residues, and to express the full B.1.351 mutations exhibited a 2.7- and 6.4-fold reduction, respectively, in neutralizing titer when compared to a pseudovirus possessing the D614G mutation alone, but retained sufficient neutralizing activity to ensure probable vaccine protection (11). However, dramatic decreases in the efficacy of three COVID-19 vaccines against the South African SARS-CoV-2 variant have been reported (14).

Recently, an immunoassay technique was used to measure 1.351 RBD binding to acute, convalescent and post-vaccination plasma IgG antibodies relative to the B1 (wild type) RBD (Edara *et al*.)(20). Antibody binding magnitude was expressed in arbitrary units relative to a standard provided by the immunoassay manufacturer. Apparently, the 1.351 RBD contained all the mutations, but it is unclear how the protein was produced. Utilizing this approach, these investigators found that antibody binding to the 1.351 RBD was 3-fold lower than to the B1 RBD, which corresponded to a 3.5-fold lower neutralizing titer. They concluded that this reduction was not sufficient to compromise the protective efficacy of the antibodies.

Next to Edara *et al*., the study reported here is one of the first to reflect direct ligand binding of a mutant RBD, rather than interference with whole virus infectivity. Contrary to expectations, we found no significant difference for the affinity of N501 and Y501 RBD binding to the ACE-2 receptor protein, and the Y501 RBD exhibited nearly twice the affinity for a human monoclonal IgG antibody specific for the wild type RBD as the N501 RBD. It has been speculated that the hydrogen bond between the Y501 residue and the Y41 residue of the ACE-2 protein contributes to an enhanced binding affinity compared to the N501 residue (9, 15, 17, 18). The O-H··N hydrogen bond, however, is stronger than the O-H··O bond, as opposed to the weaker N-H··O bond (21) and, since the N501 residue donates a hydrogen bond to the ACE-2 G352 (17), it is probably the recipient of the hydrogen bond donated by the Y41 residue. On the other hand, the N501-Y41 hydrogen bond is clearly sub-optimally aligned and separated by 3.7 Å, while the N487-Y83 bond is 2.7 Å (15), but it is unlikely that the Y501 mutant residue would be more favorably aligned.

In any case, the results of this study support the conclusion that the enhanced binding of the B.1.351 virion to the ACE-2 receptor and its decreased binding affinity for IgG antibodies elicited by the D614G pandemic strain are due to spike protein RBD conformational changes provoked by multiple mutations, rather than the effect of a single mutation. These findings may help to a) implicate the nature of how viral mutations can increase transmissibility and virulence; b) provide an experimental approach to understanding mechanisms of interference with protective immunity; and c) suggest strategies for elucidating these mechanisms.

In conclusion, a method has been developed, utilizing validated ELISAs for quantitation of human IgG antibody binding to SARS-CoV-2 spike protein and the SP receptor binding domain (RBD), to compare the binding affinities of mutant RBDs for both the ACE-2 receptor protein and human anti-SP IgG antibodies. Using this approach, we have found that the N501Y RBD mutation alone does not account for the altered ACE-2 and antibody recognition of the coronavirus variants possessing this mutation and suggests that these altered properties may be due to RBD conformational changes resulting from multiple mutations.

## ACKNOWLEDGEMENTS

This work was supported by funds from the Texas A&M University System, as part of the BADAS Study (Bacillus Calmette-Guérin Vaccination as defense against SARS-CoV-2. A Randomized Placebo-Controlled Trial to Protect Health Care Workers by Enhanced Trained Immune Responses).

